# Arbuscular mycorrhizal fungal genotype and nuclear organization as driving factors in host plant nutrient acquisition and stable carbon storage

**DOI:** 10.1101/2024.11.07.622487

**Authors:** Robert Ferguson, Ken Mugambi, Matthew Villeneuve-Laroche, Cynthia M. Kallenbach, Pedro M. Antunes, Nicolas Corradi

## Abstract

Arbuscular mycorrhizal fungi (AMF) are obligate root symbionts of most plants that improve plant growth by transferring nutrients into plant roots through networks of soil hyphae. These hyphal networks represent a carbon sink in soil; thus, it has been suggested that these fungi can also boost atmospheric carbon storage, highlighting their potential role in managing greenhouse emissions. In this study, we aimed to determine whether certain AMF genotypes and nuclear organizations (homokaryons vs heterokaryons) are associated with higher rates of host plant yield and carbon storage.
We compared Sudan-grass (*Sorghum × drummondii*) AMF inoculation across eight strains of *Rhizophagus irregularis*: four homokaryotic and four heterokaryotic strains. Sudan-grass was grown in a growth chamber, which included ^13^C-CO_2_ pulse labeling to track plant carbon into AMF.
AMF inoculation increased total and belowground biomass, as well as phosphorous, magnesium, and manganese uptake in the host. Heterokaryons led to greater belowground biomass, as well as less variable increases in shoot phosphorous. Mycorrhizal inputs to soil mineral-associated organic carbon − a highly persistent carbon pool with slow turnover − were overall greater in heterokaryons than in homokaryons but varied significantly among strains.
This indicates that the potential for carbon storage by mycorrhizal carbon inputs varies based on fungal genomic identity and nuclear organization. Overall, inoculation improved the yield of Sudan-grass and resulted in significant inter-strain variation in persistent carbon contributions to the soil. This work highlights the importance of considering genotype and nuclear identity in assessments of AMF as bio-stimulants and drivers of carbon storage.

**Societal Impact Statement:** It is crucial to develop strategies for reducing our continued excessive global increases in fertilizer applications and to offset CO_2_ emissions. The pervasive underground hyphal networks of arbuscular mycorrhizal fungi (AMF) present an enticing bio-stimulant and carbon sink. We inoculated Sudan-grass plants with eight genotypically distinct strains of a model AMF species to determine if strain identity affects plant growth and carbon storage. We found that plant biomass, nutrient acquisition and stable soil carbon inputs varied among strains, emphasizing the importance of AMF strain identity in the selection of AMF inoculants for optimizing crop yield and carbon storage.

## Introduction

Arbuscular mycorrhizal fungi (AMF) are plant root symbionts that make up the subphylum Glomeromycotina (Spatafora *et al*., 2017). These widespread organisms participate in a keystone symbiotic relationship with the roots of most terrestrial plant families called the arbuscular mycorrhizal symbiosis (Treseder & Cross, 2006). In this symbiosis, the fungal partner colonizes the plant’s roots by penetrating the epidermis, and forms highly branched tree-like structures called arbuscules in root cortical cells (Smith & Read, 2008). The arbuscules act as a trading interface, through which the fungus provides the plant host with essential nutrients in exchange for carbon (C) sources, primarily sugars and lipids (Luginbuehl *et al*., 2017). AMF are known primarily for providing the plant with phosphorous, but there is evidence that it can also increase the uptake of other essential nutrients including nitrogen, potassium, and magnesium (Chen *et al*., 2017; Fall *et al*., 2022). The response in host plant growth to mycorrhizal inoculation is referred to as the mycorrhizal growth response (MGR) (Maherali, 2014).

AMF possess a unique cellular biology, as their hyphae are coenocytic (aseptate) and can contain thousands of haploid nuclei (Kokkoris *et al*., 2020). Most strains are deemed to be homokaryotic (AMF homokaryons) (Kokkoris *et al*., 2021; Oliveira *et al*., 2024), meaning their nuclei consist of only one single haplotype, but genome and single nucleus work revealed that strains with nuclei deriving from distinct parental haplotypes also exist (AMF heterokaryons) (Ropars *et al*., 2016; Chen *et al*., 2018; Sperschneider *et al*., 2023). The presence of two distinct nuclear organizations in AMF shifts long held paradigms about their genetics and biology by demonstrating that these organisms follow both clonal and (para)sexual life stages, like more distant sexual fungi (Corradi & Brachmann, 2017). Notably, like other multinucleate fungi, the relative frequencies of the two parental haplotypes in heterokaryotic AMF can vary among strains and hosts (Kokkoris *et al*., 2021). These nuclear dynamics correlate with gene expression, with each parental haplotype acting as a distinct regulatory unit in these organisms (Sperschneider *et al*., 2023). Nuclear haplotypes can be distinguished by the identity of a putative AMF mating-type (MAT) locus (Ropars *et al*., 2016; Chen *et al*., 2018). For example, the two parental haplotype populations present in the heterokaryotic strain A4 respectively possess the MAT-1 and MAT-2 variants (Chen *et al*., 2018). This allows for direct quantification of the parental haplotype ratios within a single heterokaryotic strain using targeted DNA amplification techniques such as droplet digital PCR (ddPCR) (Kokkoris *et al*., 2021).

Each AMF nuclear organization is also associated with differing cellular and ecological traits (Koch *et al*., 2006; Ehinger *et al*., 2009). For example, AMF heterokaryons possess more nuclei than their closely related homokaryotic relatives (Kokkoris *et al*., 2021). They also differ in life history strategies, with homokaryons having faster and more successful germination, leading to higher rates of colonization, while heterokaryons grow faster after germination and produce more complex hyphal networks (Serghi *et al*., 2021). The relative abundance of co-existing nuclei in AMF heterokaryons can also change depending on the host and/or environmental conditions (Kokkoris *et al*., 2021; Cornell *et al*., 2022; Sperschneider *et al*., 2023), a phenomenon which can provide an adaptive advantage in distantly-related heterokaryotic Ascomycetes (Jinks, 1952; Zhang *et al*., 2019). This apparent distinction in fitness-associated life history strategies raises the question as to how nuclear organization may affect other AM fungal traits and plant fitness.

As organisms that grow entirely underground, AMF inhabit an ecological niche that allows them to directly participate in biogeochemical cycles (Liu *et al*., 2021). A fraction of the C fixed from the atmosphere by an AMF-colonized plant is passed to the fungus, where it may be transported throughout the mycelium or stored in vesicles and spores (Willis *et al*., 2013). Populations of AMF form extensive networks of extra-radical hyphae which can make up a considerable proportion of soil microbial biomass and, consequently, of total soil organic C (Wilson *et al*., 2009). As such, mycorrhizal exudates and necromass directly contribute inputs to soil organic carbon (SOC) and are thought to preferentially accumulate in the mineral-associated organic carbon (MAOC) pools (Ekblad *et al*., 2013; Frey, 2019; Liu *et al*., 2021). MAOC has a greater soil residence time than the larger fractions of SOC, making it an important factor when assessing C capture and the longer term persistence of accumulated fungal carbon (Emde *et al*., 2022).

AMF can further contribute to C accumulation and storage via the release of organic exudates, which slow soil C decomposition and turnover by forming stable soil aggregates (Treseder *et al*., 2007). While AMF have been found to sometimes decrease total soil C via the priming of saprophytes (Cheng *et al*., 2012; Horsch *et al*., 2023), recent evidence suggests that their contributions to SOC may be net-positive in the long-term (Verbruggen *et al*., 2013; Liu *et al*., 2021). Horsch et al. (2023) compared C inputs between AMF communities that differ in life history strategies and found that the communities associated with higher hyphal growth and turnover rates contributed most to new MAOC.

While multiple studies have also demonstrated that AMF can improve host nutrient acquisition (Smith & Read, 2008; Hodge *et al*., 2010; Luginbuehl *et al*., 2017; Ebbisa, 2023; Horsch *et al*., 2023) and atmospheric C storage (Wilson *et al*., 2009; Frey, 2019; Horsch *et al*., 2023), none have compared these factors together across multiple related fungal strains. It has become increasingly evident that the effects of AMF on soil chemistry are distinctly multifactorial (Frey, 2019). This highlights the need for further investigation into specific traits and genetics that optimize nutrient acquisition and C storage in this ubiquitous plant symbiont. In particular, the distinction in nuclear organization, germination rates and hyphal growth between AMF homokaryons and heterokaryons poses an enticing question that may help fill this knowledge gap.

The primary objectives of this study were to [1] characterize the mycorrhizal growth response (MGR) of Sudan-grass (*Sorghum x drummondi*) to eight genotypically distinct strains of *Rhizophagus irregularis*, and [2] investigate the effects of genotypic identity and nuclear organization on MGR and soil C storage from mycorrhizal inputs. We hypothesize that heterokaryons have a higher potential for C accumulation due to their adaptable nuclear dynamics, as well as their greater hyphal growth rates and mycelial network complexity, which would increase AMF-derived C inputs to the soil (Serghi *et al*., 2021; Horsch *et al*., 2023). Accordingly, we also predict that when compared to homokaryons, heterokaryons contribute relatively more to new total SOC and MAOC.

## Materials and Methods

### Inoculum Preparation

For each *Rhizophagus irregularis* strain of interest, stock root organ cultures were produced and maintained for four months to be used as inoculant for the growth chamber experiment. Strains included four homokaryotic (414, B3, C2, 330) and four heterokaryotic (A4, A5, G1, SL1) strains of *R. irregularis*. Each homokaryotic strain was selected based on its close phylogenetic relation to one of the heterokaryotic strains (**Figure S1**). Clusters of mycelia with >100 spores in pre-existing stock cultures were excised from the hyphal compartment of a split Petri plate containing *Daucus carota Agrobacterium* T-DNA transformed roots (cultivar P68) growing on M media as described in (Doner & Bécard, 1991). For each strain of interest, the two cultures with the highest spore densities were selected for use as inoculum.

### Growth Chamber Experiment

A completely randomized growth chamber experiment was conducted with 70 experimental units of Sudan-grass (*Sorghum x drummondi*), planted in tree pots and inoculated with one of the eight selected *R. irregularis* strains of interest. Additionally, AMF-free planted and AMF-free, non-planted controls were also included. Each treatment and control group consisted of seven replicate pots. Sudan-grass seeds sterilized in 50% ethanol were germinated in moist tissue paper before being planted in pairs to 5L tree pots (Stuewe and Sons Inc.) containing field soil (sandy loam), sand and turface (1:1:1). The field soil was collected from a field in Sault Ste. Marie, ON, Canada, tested for soil texture and nutrient content (sandy loam, **Table S1**) (A&L Laboratories, London, ON, Canada), and autoclaved at 120°C for 1.5 hours. Before planting, an excision of M-medium containing >100 spores of each *R. irregularis* strain was placed one inch below the soil surface, directly below the planted seedlings. Upon seedling establishment, the shortest of the two seedlings was culled, resulting in one seedling per pot. To introduce the natural microbial community from the field soil while excluding indigenous AMF, 2 mL of a soil microbial wash (filtered twice through a 20 µm sieve to remove confounding AMF species) was added to each pot after seedling establishment. The plants were grown in a Conviron PGR15 Growth Chamber at the Ontario Forest Research Institute (OFRI) in Sault Ste. Marie for 10 weeks (14h photoperiod at 700 µmol, 70% humidity, 23°C during day, 19°C during night). Deionized water was applied to each pot twice per week throughout the growth period and 80 mL of fertilizer (Miracle-Gro 24-8-16, 0.7206 g/L) was applied to each pot at two, four, and eight weeks of growth.

### ^13^C-CO_2_ Pulsing

To separate new AMF C inputs to the soil from the existing SOC, we enriched the AMF with ^13^C, using a ^13^C-CO_2_ pulse approach, to create a distinct isotopic label between the AMF and SOC. To accomplish this, after two weeks of growth, the chamber was pulsed weekly with isotopically-labeled ^13^C-CO_2_ to establish consistent ^13^C enrichment of the experimental plants throughout the growth period. Thus, ^13^C-enriched photosynthates passed to the AMF would produce an isotopic label that could be compared among soil samples. At the start of each pulse, the chamber was sealed to prevent gas exchange with the exterior. Then, a CO_2_ scrubber (Sodasorb) was activated to rapidly lower CO_2_ levels from 420 to 280 ppm. When the appropriate low CO_2_ level was reached a gas tank containing 99% ^13^C-CO_2_ (attached via tubing to the growth chamber) was opened to return the CO_2_ concentration back to 420 ppm, with a target of 33% ^13^C enrichment in the air surrounding the experimental plants. This method was used weekly for the first five pulses. To reduce costs, the four remaining weekly pulses were conducted using ^13^C-CO_2_ obtained through the reaction of 99% ^13^C-enriched Na_2_CO_3_ with H_2_SO_4_. This reaction was conducted in a sealed Erlenmeyer flask which was attached to the growth chamber through a tube (Bromand *et al*., 2001).

### Harvest and Sample Collection

At the end of the ten-week growth period, the shoots were separated from the roots (after final height measurements) and oven-dried at 60°C for >48h. Dried shoots were stored for nutrient and ^13^C analyses. Roots were washed and oven-dried at 60°C for >48h. Before drying, separate fresh root samples were stored in 1.5 mL Eppendorf tubes at −20°C for DNA extraction, and in tissue cassettes submerged in 50% ethanol for quantification of AMF colonization. Soil was homogenized and oven-dried in glass jars at 105°C for >24h then stored. After harvest and sampling we set up a soil incubation to test the longer-term persistence of the AMF C inputs and to allow time for AMF C inputs to be decomposed by the saprotrophic community added through the initial microbial wash. Subsampled soil was homogenized and placed in glass incubation jars for a four week incubation at 25°C. The jars were sealed with parafilm (to prevent rapid moisture loss) and lightly watered every two to three days. After the incubation period, the jars were oven-dried at 105°C for >24h. All oven-dried soil samples (both incubated and non-incubated) were stored for soil carbon fractioning and ^13^C analysis.

### Root Percent Colonization

Fresh roots were cleared in a 10% KOH solution and stained in a Shaeffer’s ink-vinegar solution according to Vierheilig *et al*. (1998) with a few modifications. Briefly, roots were cleared in a 90°C 10% KOH bath for three minutes, and rinsed three times with dH_2_O before staining at 90°C for four minutes. Roots were then stored in 50% ethanol at 4°C. AMF root colonization was assessed as described by (McGonigle *et al*., 1990).

### Nutrient analysis

To quantify nutrient concentrations in the Sudan-grass leaf and stem tissue at harvest, >4 g of dried homogenized shoot tissue was sent to A&L laboratories (London, ON, Canada) for analysis of nitrogen, phosphorus, potassium, magnesium, calcium, sodium, sulphur, boron, zinc, manganese, iron, copper, and aluminum. Metals were measured via acid digestion and ICP-OES (EPA 3050B/EPA 6010B). Nitrate was measured via colourimetric analysis (4500-NO_3_-F Automated Cadmium Reduction Method). Total nitrogen was measured via combustion and thermal conductivity analysis (Dumas Method, Leco). Chloride was measured via K_2_SO_4_ extraction, followed by colourimetric analysis (4500-Cl-G Mercuric Thiocyanate Flow Injection Analysis).

### Soil Carbon Fractioning

In order to isolate the mineral-associated organic matter (MAOM, granules >53 um) soil fraction, 20 g of oven-dried soil was placed in a sealable glass jar along with 80 mL of a 5 g L^-1^ sodium hexametaphosphate solution. The soil slurry was agitated sideways on a shaker for >18 h, then washed through a 53 µm sieve and oven-dried at 105°C for 24 h, leaving only the <53 µm fraction of soil containing MAOM. This process was repeated for every replicate of both at-harvest and incubated soils, for a total of 320 isolations.

### ^13^C Soil and Plant Analyses

In total, five sample types were sent for isotopic analysis: shoot tissue, pre-incubation soil, pre-incubation MAOM soil, post-incubation soil, and post-incubation MAOM soil (n=320 soil and n=73 plant samples). ∼30 mg of soil (or 4 mg of dried shoot tissue) was pulverized using a mortar and pestle and weighed into in a 9×5 mm tin capsule (Isomass Inc., SKU: D1021). Soil and plant samples were sent to the Cornell University Stable Isotope Laboratory (COIL) for analysis on a Thermo Delta V isotope ratio mass spectrometer (IRMS) interfaced to a NC2500 elemental analyzer (introduced through a Costech PN150 autosampler) to yield %C, δ^13^C vs VPDB (Vienna Pee Dee Belemnite, a primary reference scale for δ^13^C), and atom percent (AT%) of ^13^C for each sample. For each run, the effect of signal on isotopic measurement (linearity) was checked from 200 to 600 µg to define instrument response for the determination of elemental composition, using methionine as a chemical standard. An in-house standard (GNPS) was analyzed after every ten samples to ensure accuracy and precision of the instrument. Isotope corrections were performed using a two-point normalization (linear regression) for all data using two additional international standards (USGS40 and USGS41).

### Isotope Mixing Model

An isotope mixing model, as described in Horsch *et al*. (2023), was used to estimate the contribution of C from the mycorrhizal system (plant + fungus) to the soil:

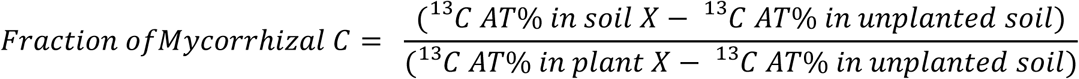

In this equation, the *Fraction of Mycorrhizal C* in soil sample X (ex: from replicate pot 7 with strain SL1) is estimated based on: the atom percent (AT%) of ^13^C in soil sample X; the AT% of ^13^C in the unplanted control soil (averaged across all control replicates for that soil type); and the AT% of ^13^C in the shoot tissues of plant A (ex: replicate plant 7 grown with strain SL1). The ^13^C AT% for each sample (all treatments and replicates, with all soil types) was run through this equation using a script in Python 3.12.3 (**Code S1**). The model output is an estimation of the fractional abundance of ^13^C contributed to the soil by the mycorrhizal system.

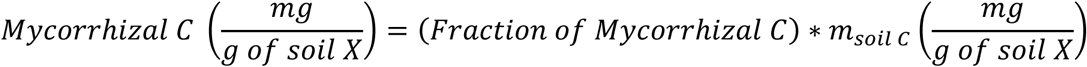

The model output was then multiplied by the total mass of C in the soil (mg g^-1^ of soil) to estimate mycorrhizal C inputs to SOC or MAOC per gram of soil.

### Droplet digital PCR

DNA was extracted from crushed roots (frozen upon harvest) from each pot containing a heterokaryotic AMF strain (A4, A5, G1, or SL1). Roots were crushed using a mortar and pestle while immersed in liquid nitrogen. 400 uL of extraction buffer (0.2 M Tris, 0.25 M NaCl, 0.025 M ethylenediaminetetraacetic acid and 0.5% SDS) was added to approximately 100 mg of crushed root, then centrifuged at 12,000 g for ten minutes. 300 µL isopropanol was added to 300 µL of isolated supernatant and centrifuged at 12000 g for ten minutes. Liquid was removed and the pellet was washed with 500 µL 70% ethanol, then centrifuged again at 12,000 g for ten minutes. The supernatant was removed, and the pellet was resuspended in EB buffer (QIAGEN). The DNA concentration was quantified (along with 260/280 and 260/230 ratios) using a NanoDrop 2000 Spectrophotometer (Thermo Scientific). Additional volumes of EB buffer were added to each tube of extracted DNA to achieve the desired DNA concentration for the PCR reaction.

PCR amplification and droplet digital PCR (ddPCR) were done on each sample to quantify the relative nuclear ratios of each pair of MAT-loci present in the heterokaryotic strains. The PCR reaction mixture in each sample consisted of 12 µL of 2x Supermix for Probes (No dUTP) (Bio-Rad, cat.no.1863023), 1.2 µL of each primers/probe mixture (Prime-Time Std qPCR assay (500 reactions), 7.2 µL of autoclaved dH20, and 1 µL of ∼50 ng/µL of extracted DNA. Each heterokaryotic strain was assigned 21 replicate wells (3 samples from the 7 replicate pots per strain) in the 96 well plate. A positive control (mycelium-extracted DNA) and a no-template control (autoclaved H_2_O) were included for each strain. Samples were emulsified using the Bio-Rad QX200 Droplet Generator with 70 µl of droplet Generation Oil for Probes per sample. 20 µL of each sample-droplet solution were transferred to a 96-well plate for PCR in a PTC Tempo Deepwell Thermal Cycler (Bio-Rad) at the following cycling conditions: 10 min at 95°C, 1 min at 94°C, 2 min at 52.5°C, followed by 44 cycles of 1 min at 94°C and 2 min at 52.2°C, then 10 min at 98°C and an infinite hold at 4°C. At every step, a ramp rate of 1°C/s was used. Following PCR amplification, ddPCR and absolute DNA quantification analysis were performed on a Bio-Rad QX200 Droplet Reader. The droplet reader identifies each droplet as positive or negative regarding the fluorescence level of the dye probe, in this case FAM or HEX. The ratio of the MAT loci within each sample was quantified using dual-target detection via multiplexing, based on the number of droplets that contained FAM and HEX fluorescence. Probes are available in **Notes S1.** The software QuantaSoft Analysis Pro (2.2.0; Bio-Rad) fits the fraction of positive droplets into a Poisson algorithm to determine the initial absolute concentration of DNA for each target signal. The positive and no-template controls were used as guides to manually set the threshold using the Threshold Line Mode Tool at the 2D amplitude mode of QuantaSoft Analysis Pro. Wells containing fewer than 10,000 droplets were excluded from the analysis. The percentage difference between the haplotypes in each strain was calculated as “((X-Y)/X)*100”, where X is the number of the most abundant haplotype and Y is the number of the least abundant haplotype.

### Statistical Analyses

All statistical tests were performed in RStudio (Version 4.3.3) (RStudio Team, 2022). We conducted one-way ANOVAs to assess the effects of AMF inoculation and nuclear organization (homokaryons vs heterokaryons), on each response variable. For analysis of the effect of AMF strain identity, a linear mixed effects model (LMM) was performed with ‘strain’ as a random factor, followed by Tukey’s HSD or Dunnett’s test for multiple comparisons. For data that could not be transformed to meet parametric assumptions, a Kruskal-Wallis non-parametric test was used, followed by Dunn’s test for multiple comparisons. Linear regressions were performed to assess effects between continuous variables. All figures were produced in RStudio.

## Results

### Effect of AMF nuclear organization and isolate genetic identity on host plant biomass

Following ten weeks in growth chambers, inoculation of Sudan-grass with *R. irregularis* strains significantly increased the host plant’s total (aboveground + belowground) biomass (average 13%, Kruskal-Wallis test, p = 0.0246) (**Figure 1A**). Although no significant difference was observed between nuclear organizations for total biomass (Dunn’s test, p = 0.8843), heterokaryons led to significantly greater belowground dry mass than the AMF-free control group (a 24% increase), while homokaryons did not (Dunn’s test, p = 0.0241, 0.1116 respectively) (**Figure 1B**). Overall, no difference was found in host belowground dry mass between heterokaryons and homokaryons when strains were grouped by nuclear organization (Dunn’s test, p = 0.4875).

**Figure 1.**
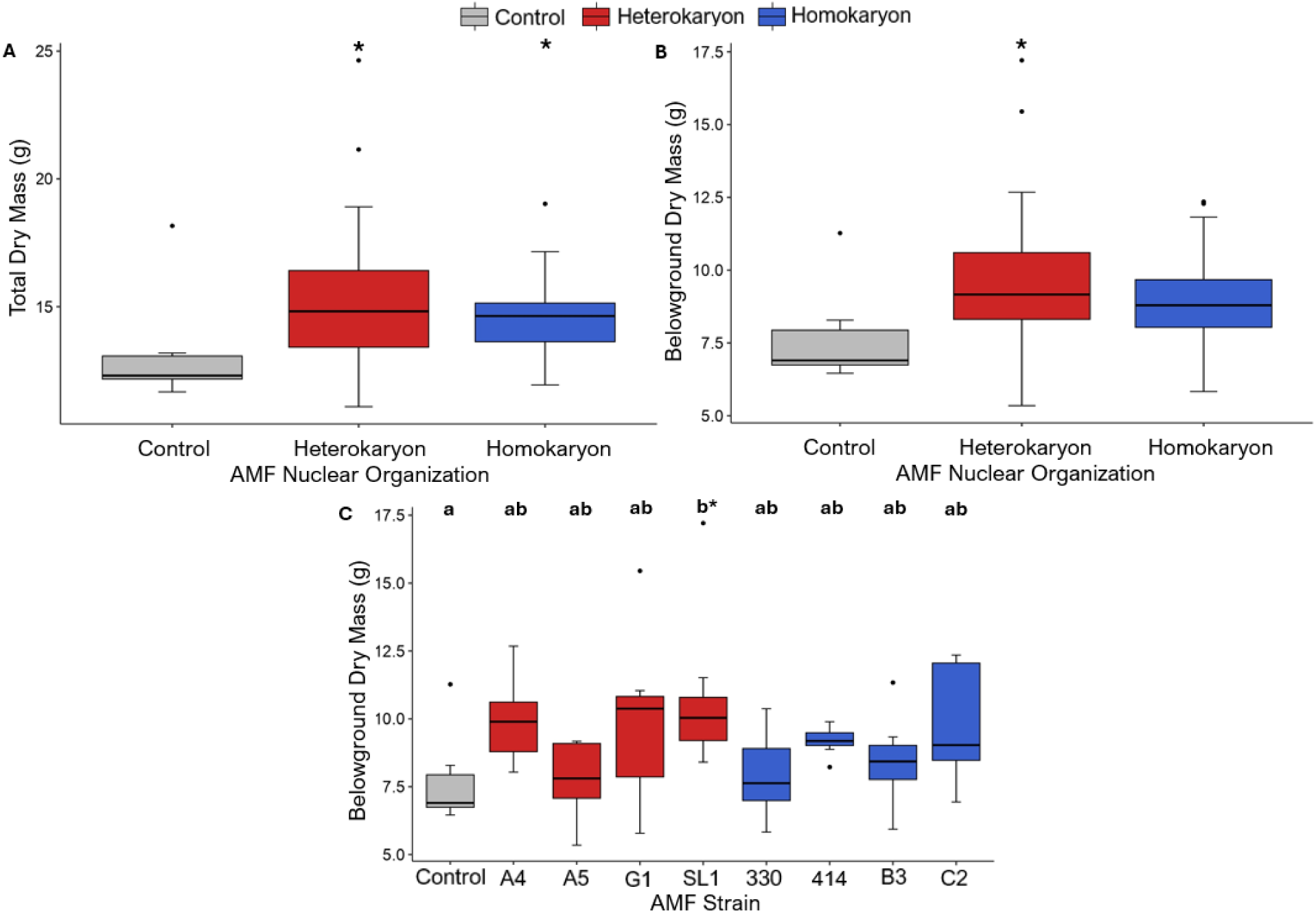
Host plant (*Sorghum x drummondii*) biomass after inoculation with one of 8 strains of *Rhizophagus irregularis*: 4 heterokaryons (in red) and 4 homokaryons (in blue), alongside a non-inoculated control (n = 63). Statistical tests were performed on log-transformed data. **A.** Average host total dry mass (g) against AMF nuclear organization (n = 7, 28, 28). **B.** Average host belowground dry mass (g) against AMF nuclear organization (n = 7, 28, 28). **C.** Average host belowground dry mass (g) against AMF strain identity (n = 7 per group). Symbols above each boxplot represent the results of Dunnett’s test and indicate a significant difference from the control group: o (0.1 > p > 0.05), * (0.05 > p > 0.01), ** (0.01 > p > 0.001), *** (p < 0.001). Letters above each boxplot represent the results of Tukey’s HSD: groups that do not share a letter are considered significantly different (p < 0.05).

When comparing individual strains, a marginal overall effect of AMF strain genetic identity on host belowground dry mass was detected (LMM, p = 0.0844) (**Figure 1C**). A Tukey’s HSD test found no significant differences among any pair of strains. The heterokaryon strain SL1 led to significantly greater host belowground biomass than the AMF-free control group (Dunnett’s test, p = 0.0378).

### Host identity and nuclear organization leading to improved nutrient uptake

Nutrient analyses revealed significant differences in Sudan-grass nutrient acquisition based on AMF strain identity. Specifically, phosphorous (P), magnesium (Mg), and manganese (Mn) were found to be significantly more elevated following inoculation, wherein differences in acquisition were linked to AMF genetic identity. For potassium, calcium, sodium, sulphur, boron, zinc, copper, and aluminum, no significant differences following inoculation were observed.

Of all nutrients investigated, host shoot P concentration was improved by an average of 65% across all strains, regardless of nuclear organization or genetic identity (ANOVA, p < 0.0001) with a significant positive effect of AMF colonization on host shoot P (linear regression, p < 0.0001) (**Figure S2A**). Specifically, shoot P was highest in the most colonized host plants, and lowest in the least-colonized host plants. There was a significant overall effect of genotypic strain identity on host shoot P concentration (LMM, p < 0.0001) (**Figure 2A**). In particular, the increases in P conferred by heterokaryons were more consistent and less variable among heterokaryon strains than the P increases among homokaryon strains. For example, while heterokaryon strains all had similar P concentrations to each other, within homokaryons, strain B3 had lower host shoot P than 414 and C2 (Tukey’s HSD test, p = 0.0269 & 0.0664). Both heterokaryons and homokaryons conferred greater host shoot P than the control (Tukey’s HSD, p < 0.0001 respectively), but were not significantly different from one another (Tukey’s HSD, p = 0.88115).

**Figure 2.**
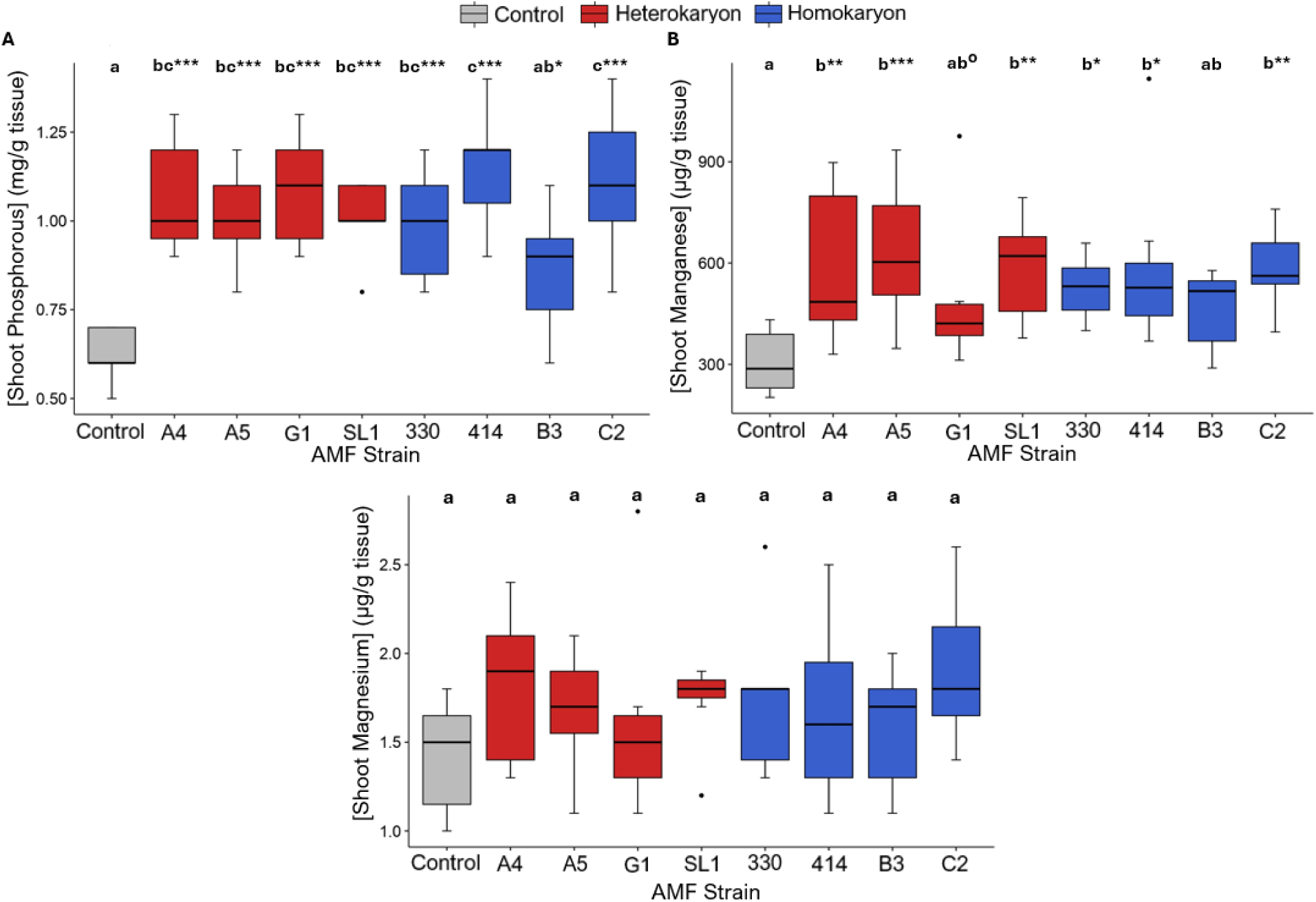
Host plant (*Sorghum x drummondii*) nutrients that were on average improved with inoculation by 8 strains of *Rhizophagus irregularis* (4 AMF heterokaryons in red, 4 AMF homokaryons in blue) alongside a non-inoculated control, against AMF strain identity (n = 63, or 7 per group). Statistical tests were performed on log-transformed data. **A.** Average host shoot phosphorous concentration (mg/g tissue). **B.** Average host shoot manganese concentration (µg/g tissue). **C.** Average host shoot magnesium (µg/g tissue). Symbols above each boxplot represent the results of Dunnett’s test and indicate a significant difference from the control group: o (0.1 > p > 0.05), * (0.05 > p > 0.01), ** (0.01 > p > 0.001), *** (p < 0.001). Letters above each boxplot represent the results of Tukey’s HSD: groups that do not share a letter are considered significantly different (p < 0.05).

When comparing all inoculated groups against the control, there was a significant improvement in Mn host shoot concentrations following inoculation (ANOVA, p < 0.0001). On average, inoculated plants had 80% greater shoot Mn than the controls. An overall effect of strain identity was also found (**Figure 2B**) (LMM, p = 0.009589). No significant differences were found among any individual pairs of strains (Tukey’s HSD, p > 0.05) nor between heterokaryons and homokaryons (Tukey’s HSD, p = 0.8791). However, all strains except for G1 and B3 led to higher shoot Mn compared to the control (Dunnett’s test, p = 0.00315, 0.00084, 0.00268, 0.01142, 0.00402, 0.00171). Mn was highest in the most well-colonized host plants, and lowest in the least-colonized host plants (linear regression, p = 0.00956) (**Figure S2B**).

Overall, AMF inoculation increased shoot Mg by an average of 21% (ANOVA, p = 0.0504), despite no overall effect of AMF strain identity (ANOVA, p = 0.492). Likewise, when grouped by nuclear organization, homokaryons and heterokaryons had similar shoot Mg (ANOVA, p = 0.149). The extent of AMF colonization had a marginally positive effect on host Mg (linear regression, p = 0.0895) (**Figure S2C**). Shoot Mg was highest in the most well-colonized host plants, and lowest in the least-colonized host plants.

### Decrease in nutrient uptake following inoculation

Inoculation of Sudan-grass led to lower acquisition of N and Fe in shoots (**Figure 3**). For example, all strains (except A4) led to reduced shoot N content (average 43%; ANOVA, p < 0.0001), with a significant effect of strain identity (LMM, p = 0.00178) (**Figure 3A**). Shoot N decreases in heterokaryons were more variable than those observed in homokaryons due to A4 conferring significantly less N reduction than G1 (Tukey’s HSD, p = 0.09579). Both heterokaryons and homokaryons as groups conferred lower shoot N than the control (Tukey’s HSD, p = 1.35×10^-4^ & 6.48×10^-5^ respectively), but were not different from one another (Tukey’s HSD, p = 0. 9404). There was a significant negative effect of AMF colonization on host shoot N (linear regression, p = 0.00578) (**Figure S3A**), with shoot N being lowest in the most well-colonized host plants, and highest in the least-colonized host plants.

**Figure 3.**
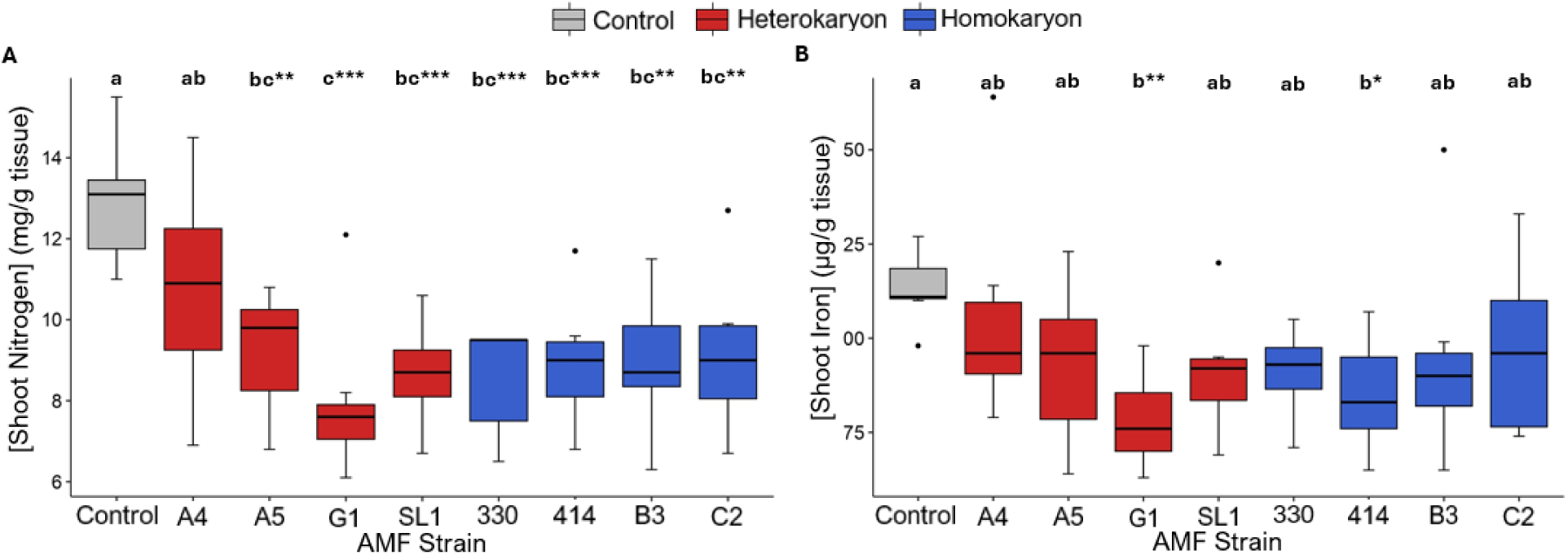
Host plant (*Sorghum x drummondii*) nutrients that were on average reduced with inoculation of eight strains of *Rhizophagus irregularis* (4 AMF heterokaryons in red, 4 AMF homokaryons in blue) alongside a non-inoculated control, against AMF strain identity (n = 63, or 7 per group). Statistical tests were conducted on log-transformed data. **A.** Average host shoot nitrogen (mg/g tissue). **B.** Average host shoot iron (µg/g tissue). Symbols above each boxplot represent the results of Dunnett’s test and indicate a significant difference from the control group: o (0.1 > p > 0.05), * (0.05 > p > 0.01), ** (0.01 > p > 0.001), *** (p < 0.001). Letters above each boxplot represent the results of Tukey’s HSD: groups that do not share a letter are considered significantly different (p < 0.05).

For Fe content, inoculation led to an average reduction of 23% (ANOVA, p = 0.00523). There was a significant effect of strain identity (ANOVA, p = 0.0436), with strains G1 and 414 leading to host plants having lower shoot Fe compared to the control (Dunnett’s test, p = 0.008, 0.05) (**Figure 3B**). Despite this, there were neither differences among any individual pair of strains (Tukey’s HSD, p > 0.05) nor between heterokaryons and homokaryons (Tukey’s HSD, p = 0.9985). There was a significant negative effect of AMF colonization on host shoot iron (linear regression, p = 0.0023) (**Figure S3B)**, and host shoot N and host shoot Fe were positively correlated (linear regression, p = 0.004771) (**Figure S4**), reflecting similar strain-specific decreases in both shoot N and Fe (**Figure 3A, 3B**).

### Soil C inputs vary depending on strain identity

While there was no significant overall effect of nuclear organization on total C inputs compared to the AMF-free control (**Figure 4A**) (Kruskal-Wallis, p = 0.8378), there was significant variation among strains in mycorrhizal inputs to MAOC after an incubation period (**Figure 4B**) (LMM, p = 0.0453). For example, the strains G1 (AMF heterokaryon) and 330 (AMF homokaryon) conferred 116% and 182% greater inputs of MAOC than strain B3 (AMF homokaryon) (Dunnett’s test, p = 0.0408, 0.0324, respectively). Significant variability in inputs existed for some strains (e.g. SL1, 330, B3) compared to others (e.g. A5, G1). AMF heterokaryons conferred marginally greater MAOC inputs than AMF homokaryons, with an average input difference of 9% (Tukey’s HSD, p = 0.0906), and there was a significant positive effect of host total dry mass on mycorrhizal C inputs to the soil (linear regression, p = 0.00469) (**Figure S5**).

**Figure 4.**
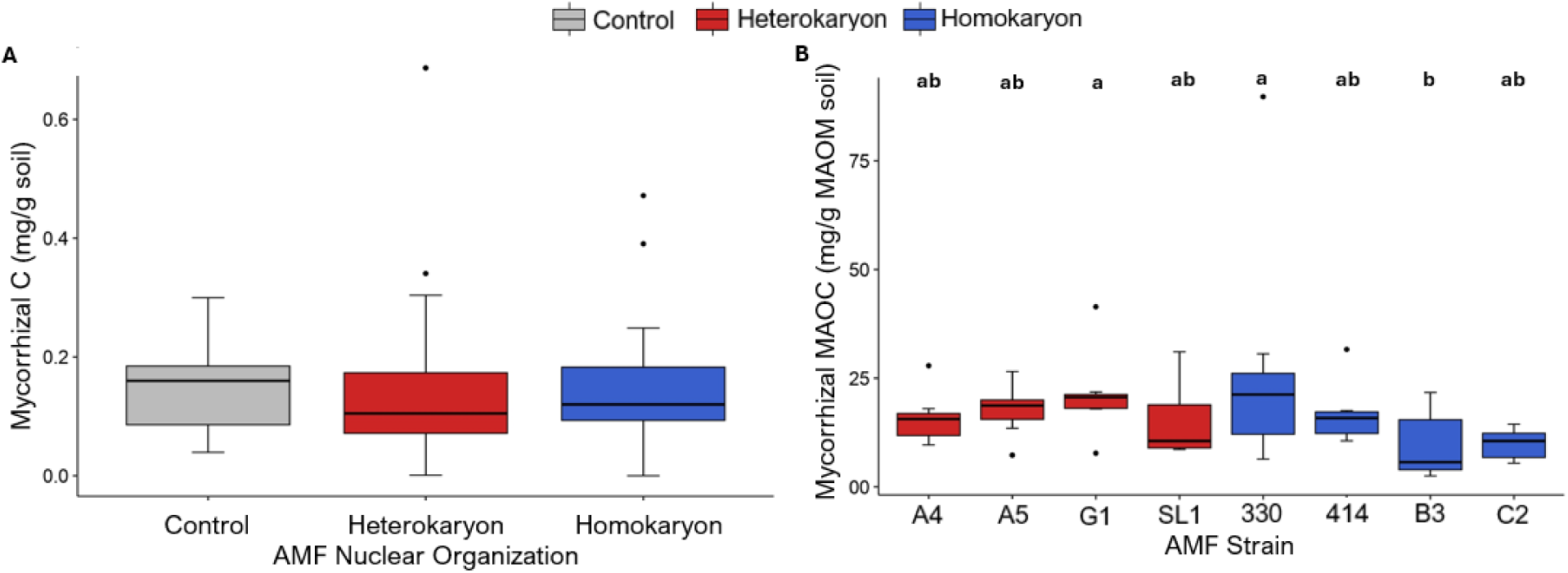
Carbon inputs contributed to the soil by the mycorrhizal system: *Sorghum x drummondi* inoculated with eight strains of *Rhizophagus irregularis* (4 AMF heterokaryons in red, 4 AMF homokaryons in blue). Statistical tests were conducted on log-transformed data. **A.** Average total carbon (C) inputs to the soil against AMF nuclear organization (n = 7, 28, 28). **B.** Average post-incubation mineral-associated organic carbon (MAOC) inputs to the soil against AMF strain identity (n = 7 per group). Letters above each boxplot represent the results of Tukey’s HSD: groups that do not share a letter are considered significantly different (p < 0.05).

### Haplotype changes in relative abundance in AMF heterokaryons with Sudan-grass

Because co-existing genomes vary in relative abundance and epigenetic contribution depending on the mycorrhizal partner, we tested for the first time the nuclear dynamics of all heterokaryons when Sudan-grass is a host. We found significant differences in the relative abundances of co-existing nuclei among AMF heterokaryons (**Figure 5**). Specifically, strain A4 carried relatively even haplotype ratios, with MAT-2 being only 17% more abundant than MAT-1. In contrast, the MAT-3 haplotype dominates in the strain A5, with 49% greater abundance compared to MAT-6, and the MAT-5 haplotype dominating in both G1 and SL1, with MAT-5 representing 75% and 23% of co-existing nuclei, respectively.

**Figure 5.**
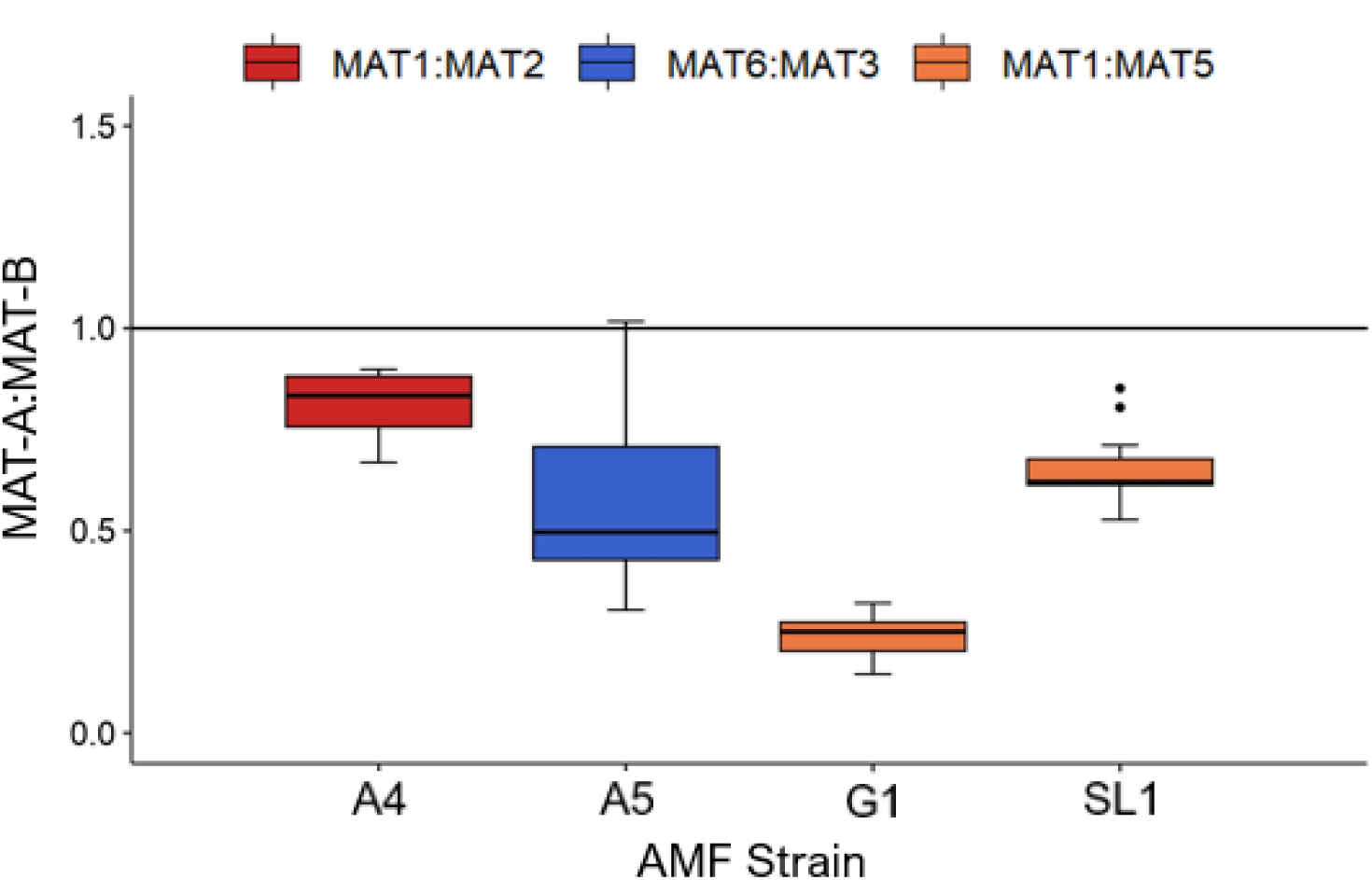
Haplotype relative abundance (MAT-A:MAT-B) in frozen root samples of each heterokaryotic strain (n = 10,18,17,13). Ratios between specific mating-type loci are denoted by colour: MAT-1:MAT-2 (A4, in red), MAT-6:MAT-3 (A5, in blue), MAT-1:MAT-5 (G1 and SL1, in orange). Data collected from droplet digital PCR analysis of DNA from crushed roots.

## Discussion

### AMF identity affects host plant biomass

Many studies have documented improved host plant performance after inoculation with AMF, including increased aboveground and belowground biomass (Qin *et al*., 2022; Chen *et al*., 2023) and plant fitness (Sohn *et al*., 2003; Campo *et al*., 2020; Terry *et al*., 2023). Notably, some experiments observe no change or negative responses in biomass after AMF inoculation (Kokkoris *et al*., 2019; Savolainen & Kytöviita, 2022; Jia *et al*., 2023), indicating that mycorrhizal colonization outcomes are dependent on mycorrhizal species, nutrient availability, and genetic background, among other factors (Koch *et al*., 2006; Schaefer *et al*., 2021; Qin *et al*., 2022; Jia *et al*., 2023; Terry *et al*., 2023).

Our results build on this notion by showing that *Rhizophagus irregularis* can significantly improve the growth of Sudan-grass, particularly in their root systems (**Figures 1A, 1B**). This supports previous studies assessing Sudan-grass in association with AMF which reported increased host aboveground and belowground biomass under normal and limiting P levels (Koyama *et al*., 2017; Kaur *et al*., 2020; Horsch *et al*., 2023).We found the degree of this improvement varied with fungal genome identity (**Figure 1C**), as post-hoc statistical comparisons found that only strain SL1 conferred a significant increase in belowground biomass over the control. There was a high degree of variation in biomass observed within the strain-specific treatments. Sudan-grass is a sorghum hybrid, and hybrid plants typically possess significant phenotypic variation (Goulet *et al*., 2017). Additionally, variation in soil conditions (Arenas *et al*., 2021), or complex interactions between the host, the AMF, and the soil microbiome (Goh *et al*., 2013) may explain some of the observed within-treatment variance. While the observed effect size of AMF strain on host biomass is small relative to that of AMF inoculation, it should be noted that this is a growth chamber study undertaken in a short period of time. Thus, the existence of a small yet significant strain effect (despite the high within-treatment variability) may propound a potential for greater variation among strains when applied at a larger scale, for a longer period of time.

The enhancement of plant growth via AMF inoculation as shown in our results is particularly relevant in the context of Sudan-grass as a cover crop grown in Ontario. Cover crops are often planted after harvest to cover agricultural soils for the purposes of reducing soil erosion, depositing organic matter, reducing nutrient leaching, improving soil fertility, reducing soil compaction, and protecting adjacent perennial crops (Scavo *et al*., 2022). Planting cover crops can thus help reduce or even reverse soil degradation that can be caused by large-scale agricultural industry practices (Spohn *et al*., 2023). Our results suggest that by conferring increases in the biomass of a common cover crop, *R. irregularis* could enhance the effectiveness of Sudan-grass in its various aforementioned roles as a cover crop, contributing to the restoration of the agricultural soil health.

### Inoculation improves phosphorous, magnesium, and manganese uptake in Sudan-grass

Phosphorous (P) is considered one of the primary nutrients influencing plant growth, which is required to maintain plant membrane structures and synthesize key metabolic macro-molecules (Malhotra *et al*., 2018). Our results showed a consistent, significant increase of phosphorous concentrations in aboveground tissues with increasing AMF root colonization (**Figure S2A**). On average, AMF-inoculated plants had close to expected phosphorous levels (0.12-0.25%), while the uninoculated plants were in the P deficiency range (A&L Laboratories Inc., 2024). The positive effect of AMF on host P uptake is well-documented in the literature (Smith & Read, 2008; Campo *et al*., 2020; Pu *et al*., 2024), and we showed this enhancement is affected by both fungal genetic identity and rates of mycorrhizal colonization (Kluber *et al*., 2012; Nguyen *et al*., 2019; Mitra *et al*., 2023).

It is noteworthy that the soil used to produce the growing medium was assessed as ‘low’ P content (**Table S1**), and no experimental plants displayed visible signs of nutrient deficiency after fertilization, meaning that P levels were not strongly limiting throughout the growth period, but significantly increased after inoculation with AMF. With P deficiency reducing yields by 30-40%, AMF inoculation thus represents a compelling, sustainable alternative to P fertilizer applications (Malhotra *et al*., 2018). In particular, all *R. irregularis* strains used in our study led to significant improvement of P acquisition across strains when grown with Sudan-grass, warranting their use to lower excessive P addition to crops in P-deficient agricultural soils.

Inoculations also improved acquisition of trace metals, such as Mg and Mn. Specifically, Mg levels post inoculation (0.17%) are close to optimal levels (0.19-0.5%) compared to levels in the deficiency range for the control (0.14%) (A&L Laboratories Inc., 2024). Mg is an essential nutrient for plant shoot and root growth, stress defence, and principally, photosynthetic performance (Hauer-Jákli & Tränkner, 2019), bearing a dominant role in photosynthetic processes in the chloroplast (Cakmak & Yazici, 2010). A meta-analysis by Hauer-Jákli & Tränkner (2019) indicates that adequate Mg supply can improve net CO_2_ assimilation by 140% and biomass by 61%, highlighting its importance for maximizing both plant yield and C storage.

For average Mn, both inoculated (558 ppm) and control groups (308 ppm) were well beyond the optimal range (5-100 ppm), indicating that the potting substrate was rich in Mn. While Sudan-grass has been found to display symptoms associated with Mn toxicity (e.g. chlorosis, necrosis, crinkling) above 200 ppm (Bowen, 1972; Li *et al*., 2019), these were not observed in this study, and the plants with the most extreme levels of Mn did not display reduced growth relative to those with the lowest levels. Under low P conditions, plants can exude organic acids such as carboxylates to solubilize inorganic P, but in doing so can chelate Mn ions in the soil, reducing the bioavailability of Mn (Hao *et al*., 2022). This may partially explain the lower Mn levels in the P-deficient control plants, alongside the fact that AMF is known to increase host Mn uptake (Fall *et al*., 2022; Sardans *et al*., 2023).

Overall, these results contribute to a large body of research demonstrating positive effects of AMF colonization on host plant Mg and Mn uptake (Cakmak & Yazici, 2010; Bücking & Kafle, 2015; Piliarová *et al*., 2019; Hauer-Jákli & Tränkner, 2019; Fall *et al*., 2022; Sardans *et al*., 2023), and the observed variation among strains provide new evidence of a significant effect of AMF genetic identity on trace metal acquisition in the host plant. Mobile trace elements are found at very low concentrations in the soil that form stable compounds that are not easily dissolved in water, and are thus less bioavailable to plants (Kaur *et al*., 2023). Despite their low concentrations, these elements are necessary for key processes, including respiration, free-radical protection and biosynthesis, making them important keystone elements for plant growth (Fall *et al*., 2022). Within this context, AMF inoculation consistently boosts their acquisition in Sudan-grass, an indication that establishment of the mycorrhizal symbiosis is of primary relevance for acquiring these crucial cellular compounds.

### Inoculation significantly reduces nitrogen and iron uptake in Sudan-grass

The role of AMF in host N nutrition has long been controversial, with studies finding negative (Savolainen & Kytöviita, 2022), neutral (Chen *et al*., 2023), and positive (Jackson *et al*., 2008; Piliarová *et al*., 2019) effects of inoculation. This effect also varies based on host identity, and those of neighboring plants (Ingraffia *et al*., 2019). For Sudan-grass, it appears that inoculation reduces N uptake, with all colonized plants being significantly below the optimal levels of 3.19-4.20% (A&L Laboratories Inc., 2024), theoretically indicating N limitations. The observed negative effect of AMF colonization on host N uptake may be the result of competition between the plant, the fungus, and other soil microorganisms for limiting bioavailable N sources (Hodge *et al*., 2010; Hao *et al*., 2022), as asserted for example in Püschel *et al*. (2016).

In mycorrhizal plants, the plant pathway (PP, via uptake transporters in the root epidermis) and fungal pathway (MP, via uptake transporters in the intra-radical mycelium and mycorrhiza-induced uptake transporters in the root cortex) present mechanisms for nutrient uptake into plant roots. With evidence that mycorrhizal colonization is capable of lowering PP activity while raising MP activity (Chiou *et al*., 2001; Harrison *et al*., 2002; Xu *et al*., 2007; Grunwald *et al*., 2009), it is possible that under N limitations the AMF will only focus on sustaining its own metabolism, while still inducing a decrease in PP N uptake. Assuming that all Sudan-grass plants were N-deficient at harvest, it is likely that our results are due to N becoming a limiting factor throughout the growth period in our experiments, leading to AMF outcompeting plants for N.

For Fe, AMF strains had an average of 92 ppm, while the control had an average of 113 ppm, both of which are within the optimal range of 54-200 ppm (A&L Laboratories Inc., 2024). This result was unexpected, as most studies find that AMF colonization tends have either positive or neutral effects on host Fe uptake (Sardans *et al*., 2023; Rajapitamahuni *et al*., 2023). Mn and Fe share similar uptake mechanisms in plant roots, and as a result, high levels of Mn can inhibit the uptake of Fe, potentially leading to deficiency (Khoshru *et al*., 2023). Considering the greater Mn levels observed in AMF-inoculated plants, this phenomenon could provide an explanation for the observed reduction of Fe levels. Moreover, the similar patterns of decreased Fe and N upon inoculation may be due to the importance of nitric oxide (NO), a key signalling molecule in Fe acquisition (Kaya *et al*., 2019; Mahawar *et al*., 2022), and indeed, limited N availability can reduce NO production, negatively affecting Fe uptake.

### AMF strain identity affects stable soil carbon inputs

Overall, our results are in agreement that AMF genetic identity can influence soil MAOC additions with Sudan-grass (Horsch *et al*., 2023). In particular, it was previously reported that the presence of members of the *Glomeraceae* family, to which the *R. irregularis* strains belong, was needed within soil communities to improve the contribution of MAOC inputs. Within this context, our analyses show that inoculation with certain AMF strains, particularly heterokaryons, can contribute to greater time-stable soil C inputs, with increased MAOC addition leading to higher potential for long term soil C storage.

The fact that MAOC inputs varied among strains while overall C inputs were similar begs the question: what mechanism could lead to the varying MAOC we observed, despite equal overall C inputs? One potential factor is the composition of the C inputs. Mycorrhizal systems that produce more MAOC-leaning compounds could add more MAOC to the soil. Specifically, MAOC primarily originates from microbial necromass and dissolved organic matter, whereas particulate organic matter, larger in size and lighter, is mainly derived from plant litter undergoing physical fragmentation and decomposition (Emde *et al*., 2022; Qi *et al*., 2023; Si *et al*., 2024). As such, variation in hyphal biomass among strains could confer variation in MAOC inputs without overall changes in C inputs from the entire system. Exudate chemical composition and concentrations also vary among different species of AMF and under differing nutrient levels (Luthfiana *et al*., 2021), along with the composition of micro-organisms associated with them (Faghihinia *et al*., 2023; He *et al*., 2023; Wen *et al*., 2023). Together, these two important biological factors could explain some of the variation in MAOC inputs between strains.

Our results also showed total C inputs from the mycorrhizal system did not differ significantly between the control and inoculated plants. This apparent lack of effect may be due to the significantly greater contribution of C from the plant (weight/C) overshadowing any differences in C inputs by the fungus. To offset this possibility, follow-up experiments should assess C storage in purely hyphal compartments (root exclusion), providing a higher resolution view of AMF-specific contribution to new soil C accumulation.

Generally, the correlation between total plant biomass and soil C inputs can be explained by the fact that greater biomass leads to higher rates of photosynthesis (Alonso-Serra, 2021), hence, the plant would have more C to pass to the soil throughout the growth period in the form of exudates and to the AMF. If these plants had been grown in the field, the C in their root systems would remain belowground even after the plant had died and a significant portion of the total biomass would be processed and sequestered below the ground in the soil (Si *et al*., 2024). Because of this, the increases in biomass conferred by AMF colonization, particularly in belowground biomass, highlights an indirect contribution to C capture whereby increases in plant growth through AMF indirectly contributes to greater C capture and storage belowground.

### AMF genome identity and nuclear organization alters plant host response and carbon storage

Our methodology, which included 4 homokaryon and 4 heterokaryon strains, provides the opportunity to assess the effects of AMF genome identity and nuclear organization on the plant host response to our factors of interest. We hypothesized that due to their adaptable nuclear dynamics, as well as their greater hyphal growth rates and network complexity, heterokaryons would have more potential for new soil C accumulation.

Remarkably, plants inoculated with AMF heterokaryons developed significantly greater belowground biomass than the AMF-free control. Expanded root systems not only benefit the plant but also improve soil quality in a variety of ways. They can reduce soil erosion, improve soil aggregation, increase water retention, and lead to increased organic matter in the soil matrix via exudates and root necromass (Piliarová *et al*., 2019). The breakdown of organic matter by microbes in the soil is an important step in nutrient cycles, by increasing levels of bioavailable nutrients for plant uptake. Increased soil organic matter can also improve soil microbial diversity, which is a key element in soil health (Bhaduri *et al*., 2022).

Overall, we found no difference in total mycorrhizal C inputs between homokaryons and heterokaryons, however, heterokaryons did confer marginally greater MAOC inputs than homokaryons. Additionally, we found that when associating with Sudan-grass, heterokaryons conferred more consistent increases in host P uptake and more variable decreases in host N uptake. Koch *et al*. (2017) found that AMF growth traits do not always directly predict host plant response. However, our hypothesis regarding the fitness advantages associated with heterokaryon strains presents a logical explanation for the observed superiority of heterokaryons in conferring increased belowground biomass and soil MAOC inputs, relative to homokaryons.

Furthermore, for nearly all our factors of interest, there was variation across our eight AMF strains, showcasing significant phenotypic differences inherent to AMF strains with distinct genetic identities. This presents an opportunity to identify specific strains that demonstrated the greatest benefits across all our results (**Figures 1-4**). The heterokaryon G1 conferred the greatest improvements in P uptake and MAOC inputs, though also demonstrated lower improvements in Mn and Mg uptake and the greatest decreases of N and Fe. The heterokaryon A4 conferred strong improvements in belowground biomass, P, Mn, Mg, and notably, the least losses in N and Fe across all strains. The homokaryon C2 demonstrated strong improvements in belowground biomass, P, Mn, and Mg, likely being the most successful homokaryon. In contrast, the homokaryon B3 demonstrated some of the weakest improvements in belowground biomass, P and MAOC inputs.

Overall, it is clear based on these observations that the advantages of specific strains are not always consistent across all variables. Thus, strain selection for inoculation with AMF should be made to produce the greatest MGR with respect to host growth and fitness. In the context of this study, the heterokaryon SL1 conferred a significant increase in host belowground biomass, making it a strong candidate for selection. It should be noted that these results pertain to the specific conditions inherent to this experiment (e.g. plant host, soil type). Future studies conducted in varied conditions could provide more exhaustive assessments of the effects of specific AMF strains on the fitness of certain host plants (e.g. seed number, reproductive biomass).

### Heterokaryon haplotype ratios with Sudan-grass differ from other plant hosts

The observed superiority of heterokaryons when grown with Sudan-grass may be due to their dynamic haplotype populations, since two co-existing haplotypes act as a distinct regulatory unit in these strains (Kokkoris *et al*., 2021; Sperschneider *et al*., 2023). This is an obvious adaptive advantage over homokaryons in these conditions, as observed in distantly-related multinucleate fungi (Jinks, 1952; Zhang *et al*., 2019).

In AMF heterokaryons, the ratios of each haplotype within a heterokaryon strain shifts dramatically in response to plant host identity, as demonstrated with multiple hosts, both in root organ cultures and greenhouse conditions (Kokkoris *et al*., 2021; Cornell *et al*., 2022; Sperschneider *et al*., 2023; Terry *et al*., 2023). Our results further support this notion, but also highlight the remarkable malleability of the AMF heterokaryotic genetic systems. Specifically, some AMF heterokaryons show ratios that are very different from those found with other hosts and conditions (Kokkoris *et al*., 2021; Cornell *et al*., 2022; Sperschneider *et al*., 2023). For example, the MAT-3 and MAT-6 nuclei in A5 have typically hovered at (or relatively close to) balance in other tested conditions, yet they deviate remarkably with Sudan-grass, differing by almost 50% on average. In contrast, the strain G1 is remarkably stable, with MAT-1 and MAT-5 nuclei always differing dramatically in relative abundance, regardless of host identity or environmental conditions (Kokkoris *et al*., 2021; Cornell *et al*., 2022).

### Conclusions

It is crucial to develop strategies for reducing our continued excessive global increases in fertilizer applications and to offset CO_2_ emissions (Lal, 2004). Our study indicates that AMF genome content and nuclear organization can influence the capacity for AMF inoculation to reduce the need for artificial nutrient addition and to contribute stable C to the soil. As such, strain selection should always be taken into consideration to improve a mycorrhizal system’s potential for plant growth and long-term C storage.

It is clear that many factors affect C storage by the mycorrhizal system, including host/fungal identity, biomass, soil nutrient content, and interactions with other microbes in the soil (Cheng *et al*., 2012; Ekblad *et al*., 2013; Parihar *et al*., 2020; Horsch *et al*., 2023). Future studies should seek to further disentangle these factors, to determine what conditions can be managed to optimize C storage in the mycorrhizal system. It will also be important to investigate the links between strain-specific growth traits and gene expression pathways, ideally using additional plant hosts, and how they affect plant response and C storage. Such research could help identify site-specific AMF-plant combinations that could inform large-scale agricultural and ecosystem restoration projects in their choice of AMF inoculation options to optimize crop yield and C storage.

## Acknowledgements

We thank the Ontario Forest Research Institute (OFRI) for providing access to facilities and to the CO_2_ climate-controlled growth chamber. We also thank D. Derbowka for his invaluable technical assistance and support throughout this research. We thank D. Storms and K. Karhi for help with harvest and sample collection, F. Hutchinson and D. Whitehead for help with soil sample processing and K. MacColl for feedback on a previous version of the manuscript. This research is funded by a MITACS Accelerate Grant to NC and PMA (IT16902). This work is also funded by the Discovery programme of the Natural Sciences and Engineering Research Council (RGPIN2020-05643 and RGPIN-2023-04103), a Discovery Accelerator Supplements programme (RGPAS-2020-00033). NC is a University of Ottawa Research Chair in Microbial Genomics and PMA is a Canada Research Chair (Tier II).

## Author Contribution

RF performed the experiments and measurements, analysed the data. KM and MVL helped with experiments and measurements. CMK provided guidance with carbon and nutrient analyses. PMA provided guidance on statistical analyses. RF and NC wrote the paper with feedback from all authors. NC and PMA designed the study and acquired funding and equipment to complete the work.

## Data Availability Statement

The raw data and R scripts are available here: https://github.com/rob-ferg/Ferguson-et-al_2024.git

