## Supplementary Figures and Tables for "Arbuscular mycorrhizal fungal genotype and nuclear organization as driving factors in host plant nutrient acquisition and stable carbon storage"

### *Plants, People, Planet* Supporting Information

The following Supporting Information is available for this article:

**Figure S1.** Phylogenetic tree of the lineages of the *R. irregularis* strains of interest.

**Figure S2.** Host plant nutrients that were on average improved with inoculation of 8 strains of *R. irregularis*, plotted against percent (%) AMF colonization.

**Figure S3.** Host plant nutrients that were on average reduced with inoculation of 8 strains of *R. irregularis*, plotted against percent (%) AMF colonization.

**Figure S4.** Linear regression of host shoot iron plotted against shoot nitrogen.

**Figure S5.** Linear regression of soil carbon inputs plotted against total dry mass.

**Table S1.** Soil test results for three field soil samples.

**Notes S1.** ddPCR Primer/Probe mixtures.

**Code S1.** Github link containing raw data and scripts.

**Figure S1.** Phylogenetic tree of the lineages of 52 *R. irregularis* strains. The strains of interest in this study are highlighted (heterokaryons in red, homokaryons in purple): A4 & 414 (DAOM229457 (MAT-6)), SL1 & 330 (T31D (MAT-2)), G1 & C2, A5 & B3. Clades are denoted as roman numerals. Mating-type (MAT) identities are denoted in parentheses. From Sperschneider et al. (2023).

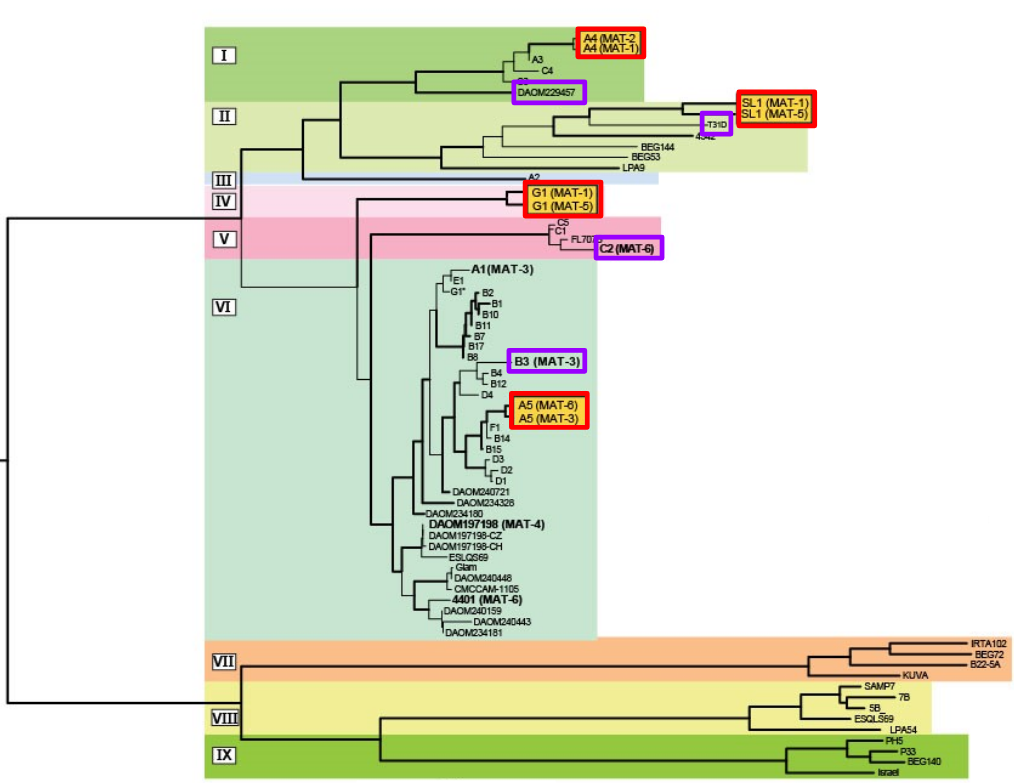

**Figure S2.** Host plant (*Sorghum x drummondii*) nutrients that were on average improved with inoculation of 8 strains of *Rhizophagus irregularis*, plotted against percent (%) AMF colonization (n = 73). A regression line is fitted to the data. A) Host shoot phosphorous concentration (mg/g tissue) (y = 0.0044337x + 0.7763, R2 = 0.2468, p = 4.66x10-6). B) The logarithm of host shoot manganese concentration (µg/g tissue) (y = 0.00480x + 6.0126, R2 = 0.078, p = 0.00956). C) The logarithm of host shoot magnesium concentration (µg/g tissue) (y = 0.002x + 0.4106, R^2^ = 0.026, p = 0.0629).

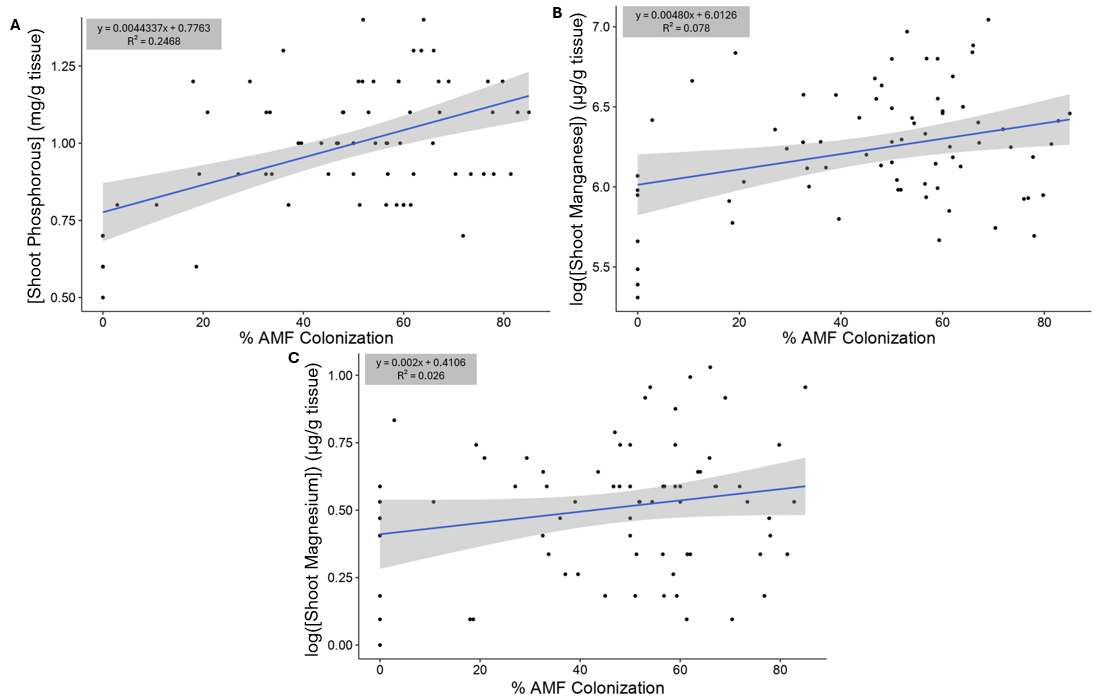

**Figure S3.** Host plant (*Sorghum x drummondii*) nutrients that were on average reduced with inoculation of 8 strains of *Rhizophagus irregularis*, plotted against percent (%) AMF colonization (n = 73). A regression line is fitted to the data. A) The logarithm of host shoot nitrogen concentration (mg/g tissue) (y = -0.003106x + 2.393803, R2 = 0.08977, p = 0.00578). B) The logarithm of host shoot iron concentration (µg/g tissue) (y = -0.003016x + 4.6622, R^2^ = 0.1111, p = 0.002301).

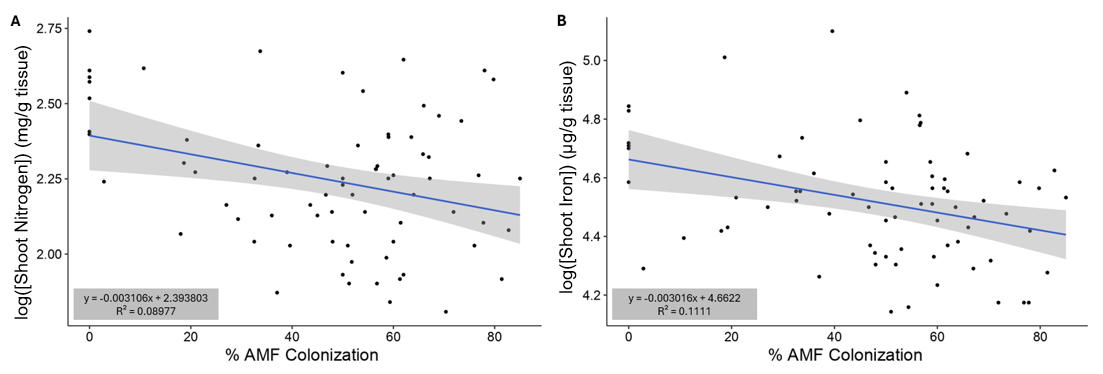

**Figure S4.** Linear regression of the logarithm of host shoot iron (µg/g soil) plotted against shoot nitrogen (mg/g tissue) of all experimental plants (*Sorghum x drummondii* inoculated with one of 8 strains of *Rhizophagus irregularis*, and a non-inoculated control group) (n = 73). A regression line is fitted to the data (y = 0.04108x + 4.13614, R^2^ = 0.1692, p = 0.004771).

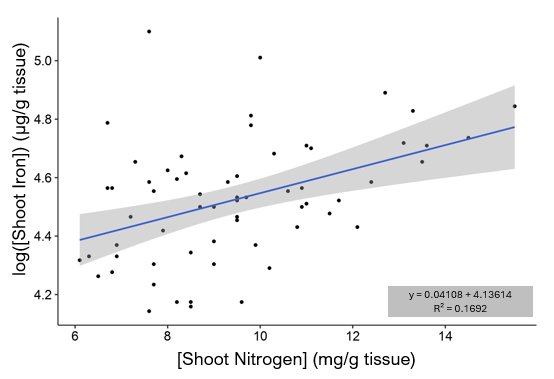

**Figure S5.** Linear regression of the logarithm of soil carbon inputs (mg/g soil) by the mycorrhizal system (*Sorghum x drummondii* inoculated with one of 8 strains of *Rhizophagus irregularis*) plotted against the total dry mass (g) of all experimental plants, including the control group (n = 73). The statistical test was performed on log-transformed data. A regression line is fitted to the data (y = 0.014956x – 0.089057, R^2^ = 0.1219, p = 0.002931).

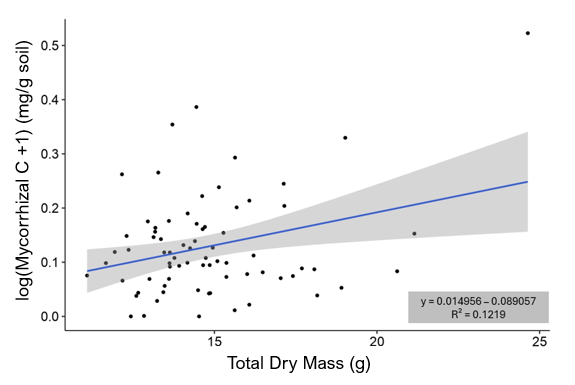

**Table S1.** Soil test results for three field soil samples (A&L Canada Laboratories Inc.).

| **Soil Sample** | **Organic Matter** | **P (ppm)** | **K (ppm)** | **Mg (ppm)** | **Ca (ppm)** | **Na (ppm)** | **Al (ppm)** | **pH** | **Buffer** | **K/Mg Ratio** |
| --- | --- | --- | --- | --- | --- | --- | --- | --- | --- | --- |
| 1 | 1.6 | 11 L | 133 M | 96 M | 330 VL | 26 H | 825 | 4.8 | 6.6 | 0.42 |
| 2 | 1.3 | 7 L | 96 M | 74 M | 260 VL | 18 H | 557 | 5.4 | 6.8 | 0.4 |
| 3 | 1.2 | 5 L | 107 M | 75 H | 260 VL | 19 VH | 534 | 5.2 | 6.9 | 0.44 |

VL = Very Low, L = Low, M = Medium, H = High, VH = Very High

**Notes S1.** ddPCR Primer/Probe mixtures.

A4: MAT-1 (Fluorescein amidite labeled, FAM) primers 5’-CATCAACAAGTCAACGATTTAT-3’ and 5’-GTGGATACATGACATGGTGT-3’ and probe 5’-CAGAAACATTTAATAATAATAATACACGTT-3’; MAT-2 (Hexachlorofluorescein labeled, HEX) primers 5’-TACACAACAAGTCAACGATG-3’ and 5’-CATGATGCTCAATATTAAGTG-3’ and probe 5’-GTAATGAAATTATAGAAGGAAATATTAG-3’.

A5: MAT-3 (HEX) primers 5’-CGTCAAAGAATCACGACACTC-3’ and 5’-CATTATTCACAATTGCGTTCGG-3’ and probe 5’-CATATAAGAAACAAAAATGCCTTAATTCAAG-3’; MAT-6 (FAM) primers 5’-CCGTCAAAGACTCACAACAC-3’ and 5’-CATTATTCACAATTGCGTTCGG-3’ and probe 5’-CACATCCGTATATATGTGCAGATAAACAA-3’.

G1: MAT-1 (FAM) primers 5’-CATCAACAAGTCAACGATTTAT-3’ and 5’-GTGGATACATGACATGGTGT-3’ and probe 5’-CAGAAACATTTAATAATAATAATACACGTT-3’; MAT-5 (HEX) primers 5’-CAGATTTAGACAAAGATATTCG-3’ and 5’-TCCTTATTACATATTTCTACA-3’ and probe 5’-CTTGTATATCAACTGTAACATATCAG-3’.

SL1: MAT-1 (FAM) primers 5’-CATCAACAAGTCAACGATTTAT-3’ and 5’-GTGGATACATGACATGGTGT-3’ and probe 5’-CAGAAACATTTAATAATAATAATACACGTT-3’; MAT-5 (HEX) primers 5’-CAGATTTAGACAAAGATATTCG-3’ and 5’-TCCTTATTACATATTTCTACA-3’ and probe 5’-CTTGTATATCAACTGTAACATATCAG-3’.

**Code S1.** Github link containing raw data and scripts.

<https://github.com/rob-ferg/Ferguson-et-al_2024.git>
